# Streamlining remote nanopore data access with *slow5curl*

**DOI:** 10.1101/2023.11.28.569128

**Authors:** Bonson Wong, James M. Ferguson, Hasindu Gamaarachchi, Ira W. Deveson

## Abstract

As adoption of nanopore sequencing technology continues to advance, the need to maintain large volumes of raw current signal data for reanalysis with updated algorithms is a growing challenge. Here we introduce *slow5curl*, a software package designed to streamline nanopore data sharing, accessibility and reanalysis. *Slow5curl* allows a user to fetch a specified read or group of reads from a raw nanopore dataset stored on a remote server, such as a public data repository, without downloading the entire file. *Slow5curl* uses an index to quickly fetch specific reads from a large dataset in SLOW5/BLOW5 format and highly parallelised data access requests to maximise download speeds. Using all public nanopore data from the Human Pangenome Reference Consortium (>22 TB), we demonstrate how *slow5curl* can be used to quickly fetch and reanalyse signal reads corresponding to a set of target genes from each individual in large cohort dataset (*n* = 91), minimising the time, egress costs, and local storage requirements for their reanalysis. We provide *slow5curl* as a free, open-source package that will reduce frictions in data sharing for the nanopore community: https://github.com/BonsonW/slow5curl

## INTRODUCTION

Nanopore sequencing has become a key pillar in the genomic technology landscape. Platform updates from Oxford Nanopore Technologies (ONT) have enabled increasingly cost-effective sequencing of large eukaryotic genomes and transcriptomes^1,2^. However, the nanopore community continues to be hampered by large data volumes and computational bottlenecks.

An ONT device measures the displacement of ionic current as a DNA or RNA molecule passes through a nanoscale protein pore. Time-series current signal data is recorded and ‘basecalled’ into sequence reads or analysed directly^1^. The available algorithms for basecalling, identification of DNA/RNA modifications and other signal-level analyses are continually evolving^3–7^. It is therefore preferable to retain ONT raw signal data for future re-analysis. However, the raw data are large – roughly ∼1 TB for a typical human genome sample at ∼30× coverage (stored in POD5 or BLOW5 format), or ∼10x larger than the corresponding basecalled reads – which imposes significant costs during storage, retrieval and reanalysis.

Cloud computing environments are increasingly popular platforms for genomics data storage and sharing. Many large, public ONT reference datasets (both existing and under construction) are hosted on cloud, including the Human Pangenome Reference Consortium (HPRC)^8^, Telomere-to-Telomere (T2T) consortium^9^, Singapore Nanopore Expression Project (SG-NEx)^10^, 1000G ONT Sequencing Consortium, NIH Center for Alzheimer’s and Related Dementias (CARD)^11^ and Genome in a Bottle Consortium (GIAB)^12^. Open access to these resources is vital for the genomics community, however, large file-sizes can make access impractical for many users. Currently, a user wishing to reanalyse a gene/transcript/region(s) of interest within a reference sample must first download the entire >1 TB dataset to their local machine or their own cloud instance, necessitating large storage capacity, a high bandwidth connection, and incurring significant egress costs (usually borne by the host). These are significant frictions for re-analysis of even a single genome/transcriptome dataset and a major barrier for large cohort datasets.

To address this challenge, we have developed *slow5curl*, a simple command line tool and underlying software library to improve remote access to nanopore signal datasets. *Slow5curl* enables a user to extract and download a specific read or set of reads (e.g. the reads corresponding to a gene of interest) from a dataset on a remote server, avoiding the need to download the entire file. *Slow5curl* uses highly parallelised data access requests to maximise speed. Here we show how *slow5curl* can facilitate targeted reanalysis of remote nanopore cohort data, effectively removing data access as a consideration.

## RESULTS

### Slow5curl basic usage

The *slow5curl* command line tool can fetch a specific read(s) from an ONT signal dataset in binary SLOW5 (BLOW5) format^13^ stored on a remote server accessible by *http/https* or *ftp* protocols (**Fig1a**). The simple BLOW5 simple file-structure, metadata header^13^ and accompanying index file, which describes the location of each read within the file, together enable efficient extraction of reads by random access pattern (**Fig1a**). The BLOW5 index may be stored remotely (either accompanying its BLOW5 file at the same URL or at another location specified by the user) or on the user’s local machine. The index is first downloaded (unless the user specifies a local index) and loaded into memory before querying the remote dataset (**Fig1a**). By default, the index will be downloaded to a temporary location and deleted by *slow5curl* after use. Alternatively, the user may retain it by specifying an option ‘*--cache’* then provide it as a local index for subsequent commands. This avoids repeated downloading of the index when making multiple successive queries.

**Fig1.**
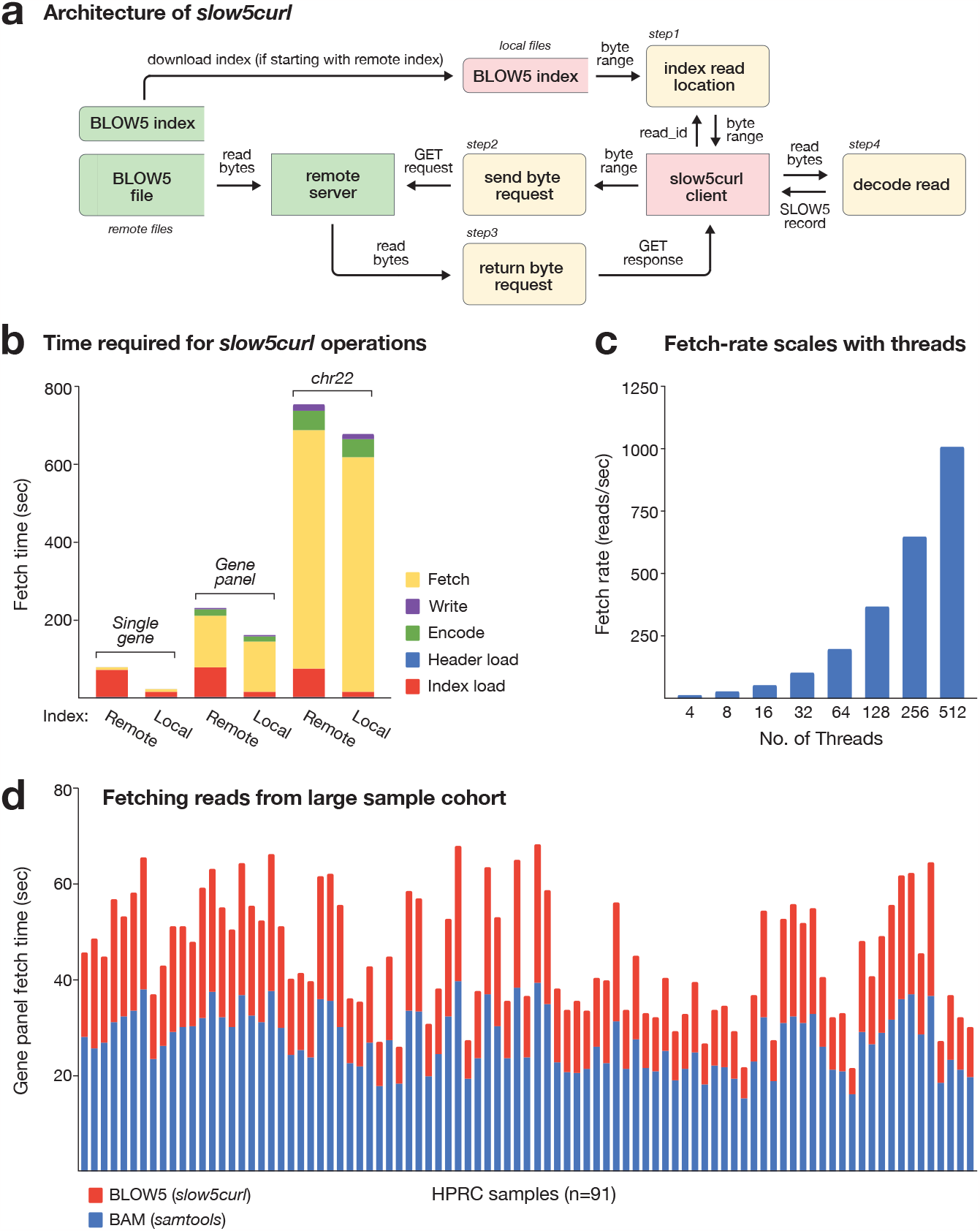
Evaluating remote nanopore data access performance with slow5curl. (**a**) Schematic summarises the data flow between entities as slow5curl fetches a single read from a nanopore signal dataset on a remote server. The slow5curl client and the remote server are represented as separate entities. Locations of datasets (BLOW5, BLOW5 Index) are denoted by their respective entity colours (green=remote; red=local). The order of execution of processes (yellow) is indicated by their accompanying step numbers. (**b**) Time taken to fetch a group of signal reads from a remote whole-genome ONT sequencing file in BLOW5 format. Times are shown separately for three sets of reads, corresponding to a single gene (left), a hypothetical gene panel comprising 100 genes (centre) and the entirety of chr22 (right). Times are shown separately for fetching reads using a remote vs local index, and overall times are broken down into the times taken for individual processes (‘fetch’, ‘write’, ‘encode’, ‘header load’, ‘index load’). Values presented are an average of n = 10 independent measurements. (**c**) Rate with which reads are fetched from the same dataset (in reads/sec) when invoking slow5curl with increasing numbers of threads (n = 4-512). (**d**) Time taken to fetch all signal reads corresponding to the hypothetical gene panel above from each of n = 91 whole-genome ONT sequencing datasets currently available via the Human Pangenome Reference Consortium (HPRC). Times are shown separately for fetching basecalled alignments (BAM format; blue) and signal reads (BLOW5 format; red) with samtools and slow5curl, respectively.

To fetch a single read or list of reads, based on their unique read IDs, the user may invoke *slow5curl get* as follows:

~~~
# get a single read with ID ‘05ef1592-a969-4eb8-b917-44ca536bec36’
slow5curl get https://url/to/reads.blow5 05ef1592-a969-4eb8-b917-44ca536bec36 -o fetched_read.blow5
# get a list of reads specified in file ‘readidlist.txt’
slow5curl get https://url/to/reads.blow5 --list readidlist.txt -o fetched_reads.blow5
~~~

In addition to *get*, the subtools *head* and *readids* may be used to print the header or a complete list of read IDs from a remote BLOW5 file, respectively.

### Fetching reads from a genomic region

A typical use-case for *slow5curl* is to fetch the raw signal reads corresponding to specific genomic region from a remote dataset. In doing so, the user may quickly re-analyse a gene/transcript of interest with the latest basecalling, DNA methylation profiling, or other signal-level analysis algorithms. Basecalled reads aligned to a reference genome/transcriptome (BAM format) must also be available, stored either locally or remotely, to provide genomic coordinates for a given read. *Slow5curl* works similarly to the remote client feature in *samtools/htslib*^14^, and the two tools may be used in tandem to retrieve raw signal reads for a specific region, as follows:

~~~
# get raw signal reads corresponding to genomic interval ‘chr1:1-1000000’
samtools view https://url/to/reads.bam chr1:1-1000000 | cut -f1 | sort -u > readidlist.txt
slow5curl get https://url/to/reads.blow5 --list readidlist.txt -o fetched_reads.blow5
~~~

To assess the performance of *slow5curl*, we measured the time taken to fetch all raw signal reads corresponding to a single gene (*BRCA1*), a hypothetical gene panel of 100 genes, or a complete chromosome (chr22) from a whole-genome ONT reference dataset hosted on our public AWS repository (https://github.com/GenTechGp/gtgseq; see **Supplementary Table 1**). Fetching reads for the single gene, gene panel and complete chromosome took 88 seconds, 254 seconds and 13 minutes, respectively, on a system with ∼3000 Mbit/s Internet connection (**Fig1b**; see **Supplementary Table 2**). Roughly ∼70 seconds was required to download the remote BLOW5 index, constituting ∼95% of the total time for the single gene. However, this was reduced to ∼13 seconds when the index was cached locally (**Fig1b**). Notably, it took ∼3.2 hours to download the whole-genome dataset using AWS Command Line Interface (AWS CLI); a significant unnecessary delay if intending to analyse only a subset.

### Efficient read-fetching by parallel threads

As shown previously^13^, BLOW5 format permits efficient parallel file access by multiple CPU threads. *Slow5curl* also uses parallel access by multiple threads to maximise performance. However, this differs from the paradigm for processor-intensive applications, wherein the ideal number of threads is close to the number of physical CPU threads available. Instead, when fetching batches of reads over the network, it is ideal to invoke an excessive number of parallel requests (e.g. hundreds) in order to hide the latency of a given request (see **Methods**).

To evaluate the multi-threading strategy used in *slow5curl*, we repeatedly fetched all chr22 reads from the ONT dataset above, each time invoking an increasing number of threads (**Fig1c**). The rate of read-fetching scaled linearly with the number of threads used and did not reach a ceiling, even with 512 threads (which was the maximum connections allowed by the server; **Fig1c**). This is indicative of highly efficient parallelisation, reducing the total time for extracting chr22 to just 294.74 seconds, of which 0.04% was loading the index (**FigS1a,b**).

### Fetching reads from a large cohort

A key motivation for developing *slow5curl* was to enable efficient access to large, public reference datasets, such as HPRC^8^. HPRC’s data is currently stored in a publicly accessible AWS bucket. Raw ONT data is stored in FAST5 format with one large tarball for each individual dataset (**Supplementary Table 3**). FAST5 tarballs do not permit indexing or random access, meaning a user must download the entire dataset for a given individual in order to access reads for even just a single gene.

To demonstrate how *slow5curl* can address this issue, we first downloaded all ONT datasets currently available from HPRC (*n* = 91), converted them to BLOW5 format with indexes (reducing the average size by 29.7%), then uploaded to commercial cloud storage (Wasabi cloud), along with accompanying basecalled alignments (see **Supplementary Methods**). From here, we used *samtools* and *slow5curl get* (as above) to remotely fetch all alignments and signal reads corresponding to our hypothetical gene panel, from each HPRC dataset (invoking *n* = 128 threads). We recorded both the time taken to fetch the reads of interest from each dataset and to re-basecall them with the latest *Guppy* version (via the *Buttery-eel* SLOW5 wrapper^15^; see **Supplementary Table 3**).

Fetching the specified reads (mean *n* = 3308 reads) from each remote file took mean 45 seconds, and a total of ∼1.2 hours was required to traverse the entire cohort (**Fig1d**). The time required for each dataset scaled linearly with their total sizes (i.e. sequencing depth), meaning the fetching rate was stably maintained across the cohort (**FigS2a,b**). Notably, the time required to basecall each set of extracted reads (mean 181 seconds) was significantly longer than its fetching time (**FigS2c**; **Supplementary Table 3**). Since basecalling can be initiated on each individual set of reads without waiting for the subsequent set to be fetched, the overall time taken to complete this analysis is almost entirely determined by the basecalling time, and the net time added for data access with *slow5curl* becomes negligible. Similarly, the experiment would require downloading ∼22.5 TB of BLOW5 files to local storage, compared to ∼120.5 GB of reads fetched by *slow5curl*, dramatically reducing data egress costs incurred on most commercial cloud platforms. Availability of such large local storage capacity is also unrealistic for most users. In summary, this experiment demonstrates how *slow5curl* can be used to dramatically reduce the overheads for data access during reanalysis of ONT cohort data.

## DISCUSSION

Data accessibility is critically important to the genomics community and a prerequisite for open, reproducible science. With the breadth of nanopore sequencing adoption and the scale of nanopore datasets growing rapidly, there is a need for new and efficient methods for nanopore data sharing and public access. *Slow5curl* allows a user to quickly fetch specific reads (e.g. for a gene of interest) from a raw nanopore signal dataset on a remote server, without downloading the entire dataset. This saves time, egress costs, and reduces the need for a high-bandwidth connection and large local storage. *Slow5curl* makes it feasible for even low-resource users to fetch and reanalyse nanopore signal data from large cohort datasets like HPRC and, in doing so, increases the value of such initiatives.

The large size and complex file structure of ONT native signal datasets poses a particular challenge for genomics data repositories, such as EBI’s European Nucleotide Archive (ENA) or NCBI’s Sequence Read Archive (SRA). ONT’s FAST5 format is currently supported by ENA and SRA. However, users must upload a single FAST5 tarball for a given datasets, which is typically >1-2 TB for a standard PromethION sequencing run. A user wishing to access the data must then download and extract the entire file. Given these barriers, many nanopore users neglect to provide the raw data for published studies to SRA or other repositories, preventing re-analysis with updated basecalling, methylation profiling or other signal-based analysis methods^3–7^. *Slow5curl* provides an improved solution for data repositories, analogous to the familiar *htslib*/*samtools* and *fqidx*/*faidx curl* protocols, which facilitate access to remote BAM and FASTQ data, respectively^14^. We anticipate that streamlined accessibility would encourage more users to share raw nanopore datasets on permanent public repositories.

In fetching specific reads from a remote dataset with minimal delay, *slow5curl* has the potential to enable interactive analysis and exploration of large nanopore signal datasets. For example, one can envision an interactive browser for signal data exploration, analogous to existing genome browsers that work with sequence-level data. While there are several current tools for visualising nanopore signal reads, such as our own recent package *Squigualiser* (https://github.com/hiruna72/squigualiser), these offer static visualisation only, and require the dataset(s) under inspection to be stored locally, which is problematic for large nanopore datasets. *Slow5curl* provides a mechanism for interactive exploration of remote data, with reads being rapidly fetched, processed and plotted as the user navigates the hypothetical browser. We show here that a cached local index would reduce the latency on this process to a matter of seconds. Further speed-ups are likely possible by integrating more specialised protocols, such as the S3 API, into *slow5curl*, although this would necessitate tradeoffs in compatibility. We chose to use the standard *curl* library for its compatibility with any *http*/*https* or *ftp* hosted storage.

*Slow5curl* is the latest feature in the SLOW5 data ecosystem, a community-centric project designed to improve the usability of nanopore signal data (https://hasindu2008.github.io/slow5/). The SLOW5 data format^13^ is now accompanied by software libraries in C/C++, python, rust and R for reading/writing SLOW5 files (https://github.com/hasindu2008/slow5lib); the *slow5tools* package for creating, converting, handling and interacting with SLOW5/BLOW5 files^16^; the *Buttery-eel* wrapper for ONT basecalling and methylation calling software^15^; the *Squigulator* and *Squigualiser* packages for simulation and visualisation of signal data^17^; and a range of other open source tools^7,18–22^. Efficient, remote data access by *slow5tools* is possible thanks to the simple SLOW5/BLOW5 file structure and accompanying index. Complex file formats like ONT’s original FAST5 or new native POD5 format do not support efficient random access or indexing, thereby prohibiting efficient remote data access. We provide *slow5curl* as a free and open resource to improve data accessibility for the nanopore community: https://github.com/BonsonW/slow5curl

## METHODS & IMPLEMENTATION

### Architecture and Implementation of slow5curl library (slow5curllib)

The underlying library *slow5curllib* is written in C; it utilises the file format library *slow5lib*, and the multiprotocol file transfer library *libcurl*. Minimising dependencies is a central design principle of the SLOW5 ecosystem. We therefore chose to develop *slow5curllib* as a separate library, rather than incorporating it into *slow5lib* or *slow5tools*, to avoid adding *libcurl* as a new dependency to these core SLOW5 packages.

Every SLOW5/BLOW5 file can be represented with a (much smaller) corresponding index file that maps every read ID to its respective location in memory. Since most RESTful APIs allow for byte-range fetches, *slow5curllib* takes advantage of this index file to send read-specific file transfers.

The library implements a single fetch (*s5curl_get()*) through the interface of *libcurl*. Once the BLOW5 header and its index is downloaded, we supply a connection handle to *libcurl* containing all the necessary configurations required to generate a byte-range request to the remote server. The thread making the call then waits until this request is fulfilled. *Slow5curl’s* batch fetch uses this exact method internally on parallel threads.

Batch fetches are a high-level multithreaded option for getting lists of reads quickly (using *s5curl_get_batch()*). *Slow5curllib* does this by spawning worker threads (C/C++ POSIX) to fetch reads in parallel. This way we can accelerate high-volume fetch operations on multi-threaded systems.

In very rare instances, for network-related reasons, one or more fetches within a batch will fail. Instead of aborting the method (since the library does not expose each worker thread), *slow5curllib* provides the option to retry any particular fetch a certain amount of times before it fails (default 1). Since it is usually an external issue, we also provide a parameter to control the amount of time to wait before retrying (default 1 sec). If a fetch fails twice in a row, it is likely that something has gone wrong with the server/connection, or the client is being denied further access.

### Architecture and Implementation of slow5curl tool

*Slow5curl* provides the functionality of the library through a command-line interface. Each *slow5curl get* command simply invokes the library method *s5curl_get()* unless provided with a list, where it will instead invoke *s5curl_get_batch()*. Additionally, *slow5curl* is able to provide BLOW5 file meta-data to the user. The *slow5curl head* command prints out the header downloaded from the remote BLOW5, and *slow5curl reads* prints out all read IDs stored in the BLOW5 index.

By default, *slow5curl* will automatically delete any downloaded BLOW5 index unless a permanent file path is specified through the --*cache* option. This option is for if the user requires to fetch data from a remote BLOW5 more than once. Downloading the index takes a non-negligible amount of time, so caching it to a local path will avoid repeated downloads. After the index is cached, the user can provide a local index path through the --*index* option.

### Benchmark experiments

#### Datasets

The HG002 (NA24385) reference dataset used for the benchmarking (**Supplementary Table 1**) was prepared using the ONT LSK114 ligation library kit and was sequenced on an ONT PromethION on an R10.4.1 flow cell to generate ∼30X genome coverage. Sheared DNA libraries (∼17Kb) were used. The FAST5 files were live-converted using the *real-f2s* script and then merged into a single BLOW5 (*zlib+svb-zd* compression) file and indexed using *slow5tools*^16^. Basecalling was performed using *Buttery-eel* (through Guppy v6.4.2) under the high-accuracy model. Reads were mapped to the hg38 reference using *Minimap2* (v2.17), and a sorted BAM file (with index) was created using *samtools*. The data was uploaded to the *gtgseq* AWS S3 bucket in the *US West (Oregon) us-west-2* region using AWS CLI.

**Supplementary Table 1.** Data specifications.

| Dataset | Platform | Pore | Sampling rate | Reads | Total seq. | Read length (mean, median, max) | BLOW5 file size (zlib+svb-zd) | BLOW5 index size |
| --- | --- | --- | --- | --- | --- | --- | --- | --- |
| ~30X human genome (NA24385) | PromethION | 10.4.1 | 4 kHz | 15.3M | 102.2 Gbases | 6.5, 7.3, 636.6 kbases | 1.1 TB | 836 MB |

The Human Pangenome Reference Consortium (HPRC) data (n=91 samples) was downloaded from the human-pangenomics AWS S3 bucket. For each sample, the downloaded tarball of FAST5 files was extracted and then was converted into a merged BLOW5 file (zlib+svb-zd compression) and indexed using *slow5tools*. The 31.2 TB of FAST5 tarballs, reduced to 21.93 TB after the BLOW5 conversion (see **Supplementary Table 3**). The available basecalled data for each sample was also downloaded (FASTQ.gz format) from the human-pangenomics AWS S3 bucket and were mapped to the hg38 genome using *Minimap2* (v2.17), then sorted and indexed using *samtools*. The BLOW5 files (with index) and BAM files (with index) for all the 91 samples were uploaded to an s3 bucket in the Wasabi cloud under the *Asia Pacific (Sydney) ap-southeast-2 region* using AWS CLI.

#### System information

A Dell PowerEdge C4140 server computer with a 10Gb ethernet network connection was used for the experiments (**Supplementary Table 2**). The server is located in Sydney and was measured to have ∼3Gbit/s download speed when benchmarked via *speedtest* by *ookla*.

**Supplementary Table 2.** Computer and connectivity specifications.

| CPU (No. of cores) | RAM | OS | Ethernet connection | Computer location | Internet connection (*speedtest*) |
| --- | --- | --- | --- | --- | --- |
| Intel Xeon Silver 4114 CPU @ 2.20GHz (20 cores/40 threads) | 384 GB | Ubuntu 18.04 LTS | Intel Ethernet Controller X710 for 10GbE SFP+ | Sydney | Download: 3405.53 Mbit/s<br>Upload: 2871.61 Mbit/s<br>Ping: 1ms<br>ISP: University of New South Wales<br>Server: AARNet - Sydney |

#### Methodology for HG002 experiments

The HG002 dataset is hosted on the AWS S3 bucket in the *US West (Oregon) us-west-2* region, and represents a high-latency scenario when being accessed from a computer located in Sydney.

We tested the performance impact of the number of reads fetched by *slow5curl* by providing read IDs corresponding to the region of *BRCA1* gene (chr17:43044295-43170245), a hypothetical gene panel comprising 100 randomly selected genes, and *chr22* (the smallest human autosome). Each test was run on 128 threads, with the average time recorded from 10 runs. All runs were performed during low-network load conditions (on weekends).

#### Methodology for HPRC cohort experiments

This dataset is stored on the *Asia Pacific (Sydney) ap-southeast-2* region, and represents a low-latency scenario when being accessed from a computer located in Sydney.

We test *slow5curl* on 91 samples alongside *samtools* to fetch all reads corresponding to a hypothetical gene panel comprising 100 randomly selected genes. This involves first using *samtools* to fetch the read IDs corresponding to the gene panel regions (BED format) into a read ID list. After this, we use *slow5curl* to fetch the reads into a BLOW5 file. Lastly, we basecall the reads using *Buttery-eel* (through *Guppy* v6.4.2) with the super-accuracy (SUP) model. This experiment was run during low network load conditions.

## DATA & CODE AVAILABILITY

The HG002 dataset in BLOW5 format used for benchmarking is available as part of the AWS Open Data Program (https://registry.opendata.aws/gtgseq/) in the *gtgseq* S3 bucket (https://gtgseq.s3.amazonaws.com/index.html). This dataset is also available under the NCBI Sequence Read Archive (SRA) at **Bioproject PRJNA744329**. HPRC data is available in FAST5 format under the human-pangenomics AWS S3 bucket (https://s3-us-west-2.amazonaws.com/human-pangenomics/index.html) and can be converted to BLOW5 format by following instructions in the **Supplementary Methods** section.

*Slow5curl* is free and open source and can be accessed at: https://github.com/BonsonW/slow5curl. The GitHub commit used for the benchmarks is 6d930a3a6cc3e206fbfc21c402a8fc59717cacfc.

## ACKNOWLEDGEMENTS

We thank the AWS Open Data Sponsorship Program for generously hosting an open dataset in BLOW5 format that greatly assisted in implementing and testing *slow5curl*. We acknowledge the following funding support: Australian Medical Research Futures Fund grants MRF1173594, MRF2016008 and MRF2023126 (to I.W.D.) and Australian Research Council DECRA Fellowship DE230100178 (to H.G.).

## DECLARATIONS

I.W.D. manages a fee-for-service sequencing facility at the Garvan Institute of Medical Research that is a customer of Oxford Nanopore Technologies but has no further financial relationship. H.G., J.M.F. and I.W.D. have previously received travel and accommodation expenses from Oxford Nanopore Technologies. The authors declare no other competing financial or non-financial interests.

## CONTRIBUTIONS

All authors (B.W., J.M.F., H.S. & I.W.D.) contributed to the conception, design and benchmarking of *slow5curl*. B.W., & H.G. implemented *slow5curl*. B.W. performed benchmarking experiments. B.W., H.G. & I.W.D prepared the figures and manuscript.

## SUPPLEMENTARY METHODS

### Commands used for benchmark experiments on HG002 dataset

#### Preparation of data

~~~
# merging and indexing live-converted BLOW5
slow5tools merge tmp_slow5/ -o PGXX22394_reads.blow5 -t40
slow5tools index PGXX22394_reads.blow5
# basecalling
buttery-eel -g ont-guppy-6.4.2/bin/ --config dna_r10.4.1_e8.2_400bps_hac_prom.cfg --device ‘cuda:all’ -i
PGXX22394_reads.blow5 -o hg2_guppy_6.4.2_hac.fastq --qscore 9 --port 5555 --use_tcp
# mapping
minimap2 -ax map-ont -t16 --secondary=no hg38noAlt.idx hg2_guppy_6.4.2_hac.pass.fastq | samtools sort - >
hg2_guppy_6.4.2_hac_pass_minimap2.17.bam && samtools index hg2_guppy_6.4.2_hac_pass_minimap2.17.bam
# upload
aws s3 cp ${FILE} s3://gtgseq/ont-r10/NA24385/${PATH} --profile gtg-open-data
~~~

#### Fetching regions with *slow5curl*

~~~
# generating the read ID list for one gene
samtools view hg2_guppy_6.4.2_hac_pass_minimap2.17.bam chr17:43,044,295-43,170,245 | cut -f 1 | sort -u >
PGXX22394_one_gene_readid.list
# generating the read ID list for hundred genes
samtools view hg2_guppy_6.4.2_hac_pass_minimap2.17.bam -L hundred_gene.random.hg38.bed -M | cut -f 1 |
sort -u > PGXX22394_hundred_gene_readid.list
# generating the read ID list for chr22
samtools view hg2_guppy_6.4.2_hac_pass_minimap2.17.bam chr22 | cut -f 1 | sort -u >
PGXX22394_reads_chr22_readid.list
# slow5curl remote index
/usr/bin/time -v ./slow5curl get https://gtgseq.s3.amazonaws.com/ont-r10-
dna/NA24385/raw/PGXX22394_reads.blow5 -t 128 --list ${READID_LIST} -o reads.blow5
# slow5curl local index
/usr/bin/time -v ./slow5curl get https://gtgseq.s3.amazonaws.com/ont-r10-
dna/NA24385/raw/PGXX22394_reads.blow5 -t 128 --list ${READID_LIST} -o reads.blow5 --index
PGXX22394_reads.blow5.idx
~~~

#### Performance scaling with number of threads when using *slow5curl*

~~~
# generating the read ID list
samtools view hg2_guppy_6.4.2_hac_pass_minimap2.17.bam chr22 | cut -f 1 | sort -u >
PGXX22394_reads_chr22_readid.list
# time slow5curl
/usr/bin/time -v ./slow5curl get https://gtgseq.s3.amazonaws.com/ont-r10-
dna/NA24385/raw/PGXX22394_reads.blow5 -t ${THREADS} --list PGXX22394_reads_chr22_readid.list -o
reads.blow5 --index PGXX22394_reads.blow5.idx
~~~

#### Downloading the whole file

~~~
# download file through AWS CLI
/usr/bin/time -v aws s3 --no-sign-request cp s3://gtgseq/ont-r10-dna/NA24385/raw/PGXX22394_reads.blow5 .
~~~

### Commands used for benchmark experiments on human pangenome reference dataset

#### Preparation of data

~~~
# FAST5 tarball conversion
tar xf ${NAME}.fast5.tar -C fast5_tmp/
slow5tools f2s fast5_tmp/ -d slow5_tmp -p40 –retain
slow5tools merge slow5_tmp/ -o ${NAME}.blow5 -t40
slow5tools index ${NAME}.blow5
# mapping
minimap2 -ax map-ont hg38noAlt.idx ${NAME}.fastq.gz -t 16 --secondary=no | samtools sort - -o ${NAME}.bam
&& samtools index ${NAME}.bam
# upload
aws s3 --profile wasabi --endpoint-url=https://s3.ap-southeast-2.wasabisys.com cp ${FILE}
s3://slow5curl/${FILE}
~~~

#### slow5curl

~~~
# generating the read ID list
/usr/bin/time -v samtools view https://s3.ap-southeast-2.wasabisys.com/slow5curl/$NAME.bam –L
hundred_gene.random.hg38.bed -M | cut -f1 | sort -u > ${NAME}_rids.list
# time slow5curl
/usr/bin/time -v slow5curl get https://s3.ap-southeast-2.wasabisys.com/slow5curl/$NAME.blow5 -t 128 --
list ${NAME}_rids.list -o ${NAME}_reads.blow5
# time basecalling
/usr/bin/time -v buttery-eel -g ont-guppy-6.5.7/bin/ -x cuda:all --config dna_r9.4.1_450bps_sup_prom.cfg
-i “out/${NAME}_reads.blow5” -o “out/${NAME}_reads.fastq”
~~~

### *Slow5curl* C library Usage

The *slow5curl* C API is built on the libraries *slow5lib* and *libcurl*. The API is split into methods for the initialisation/cleanup of: remote BLOW5 files, BLOW5 indexes, connection handles; and performing single/batch read fetches. The entire process of fetching reads is structured as so:

1. Intialise global resources
2. Initialise remote BLOW5 file
3. Download the BLOW5 index file if it is not available locally
4. Initialise the BLOW5 index
5. Initialise connection handle(s)
6. Perform single/batch fetches
7. Do something with the returned fetch
8. Cleanup initialised memory and resources

All interactions with the API should be after global resources initialised and before they are freed:

~~~
s5curl_global_init();
// … slow5curl operations
s5curl_global_cleanup();
~~~

Initialising resources required to fetch from a remote BLOW5 file is done simply by “opening” it from the provided URL and then “loading” its index:

~~~
// open remote file and load its index
s5curl_t *s5c = s5curl_open(“https://url/to/reads.blow5“);
s5curl_idx_load(s5c);
~~~

Fetching a single record requires a connection handle. This is exposed to give developers the flexibility of reusing connection handles and hence the ability to implement their own performance optimisations:

~~~
// create a connection handle for individual fetches
S5CURLCONN *conn_handle = s5curl_conn_init();
// perform a single fetch slow5_rec_t *record = NULL;
s5curl_get(s5c, conn_handle, read_id, &record);
~~~

The library also offers a method to fetch reads on multiple threads. Here connection handles reside in the core data structure and records are returned in their respective data batch:

~~~
// initialise multiple connection handles for batch fetches
s5curl_mt_t *core = s5curl_init_mt(num_threads, s5c);
// perform a batch fetch
slow5_batch_t *db = slow5_init_batch(BATCH_CAPACITY);
s5curl_get_batch(core, db, read_ids, num_reads);
~~~

Both *slow5curl* and *slow5lib* share the same return data structures (*slow5_rec_t* and *slow5_batch_t*), and hence, should offer some degree of interoperability between the two libraries.

## SUPPLEMENTARY FIGURE LEGENDS

**FigS1.**
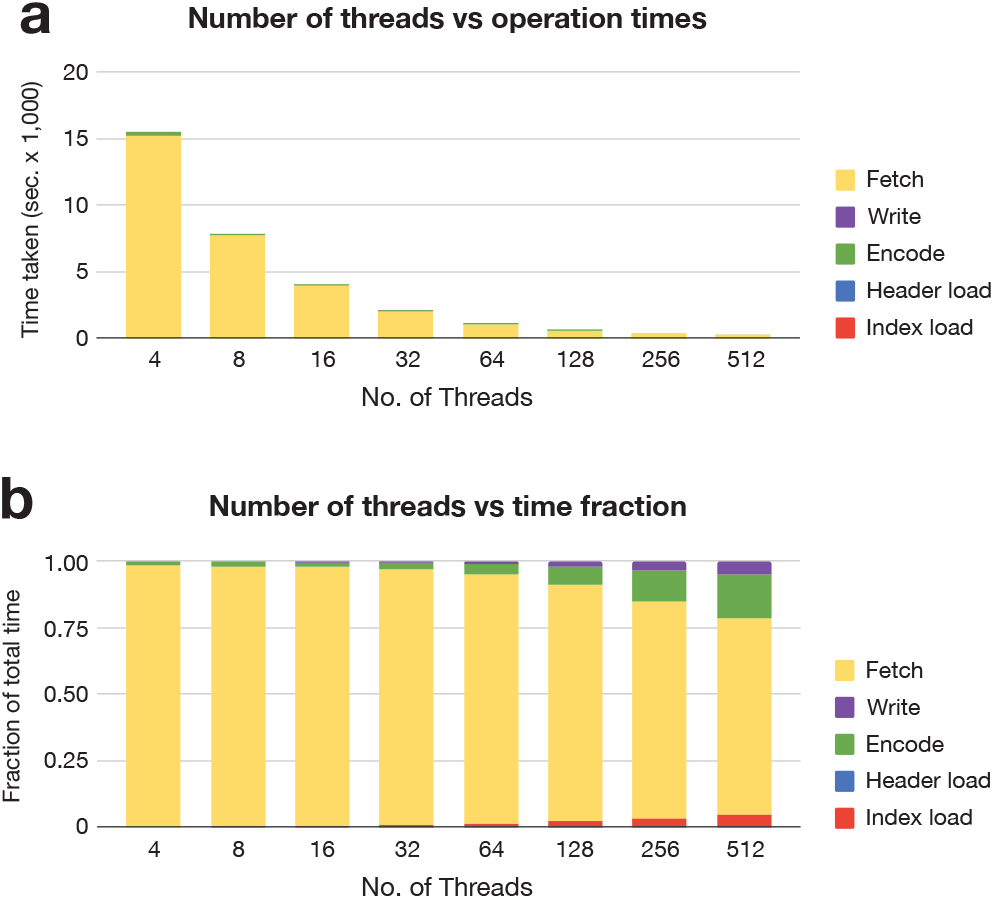
Evaluating the multi-threading performance in slow5curl. (**a**) Time taken to fetch all reads corresponding to a hypothetical gene panel comprising 100 genes from a remote whole-genome ONT sequencing file in BLOW5 format, when invoking slow5curl with increasing numbers of threads (n = 4-512). Overall times are broken down into the times taken for individual processes (‘fetch’, ‘write’, ‘encode’, ‘header load’, ‘index load’). (**b**) Same as above but times for each individual process are expressed as a fraction of the total run-time, in stacked bar format.

**FigS2.**
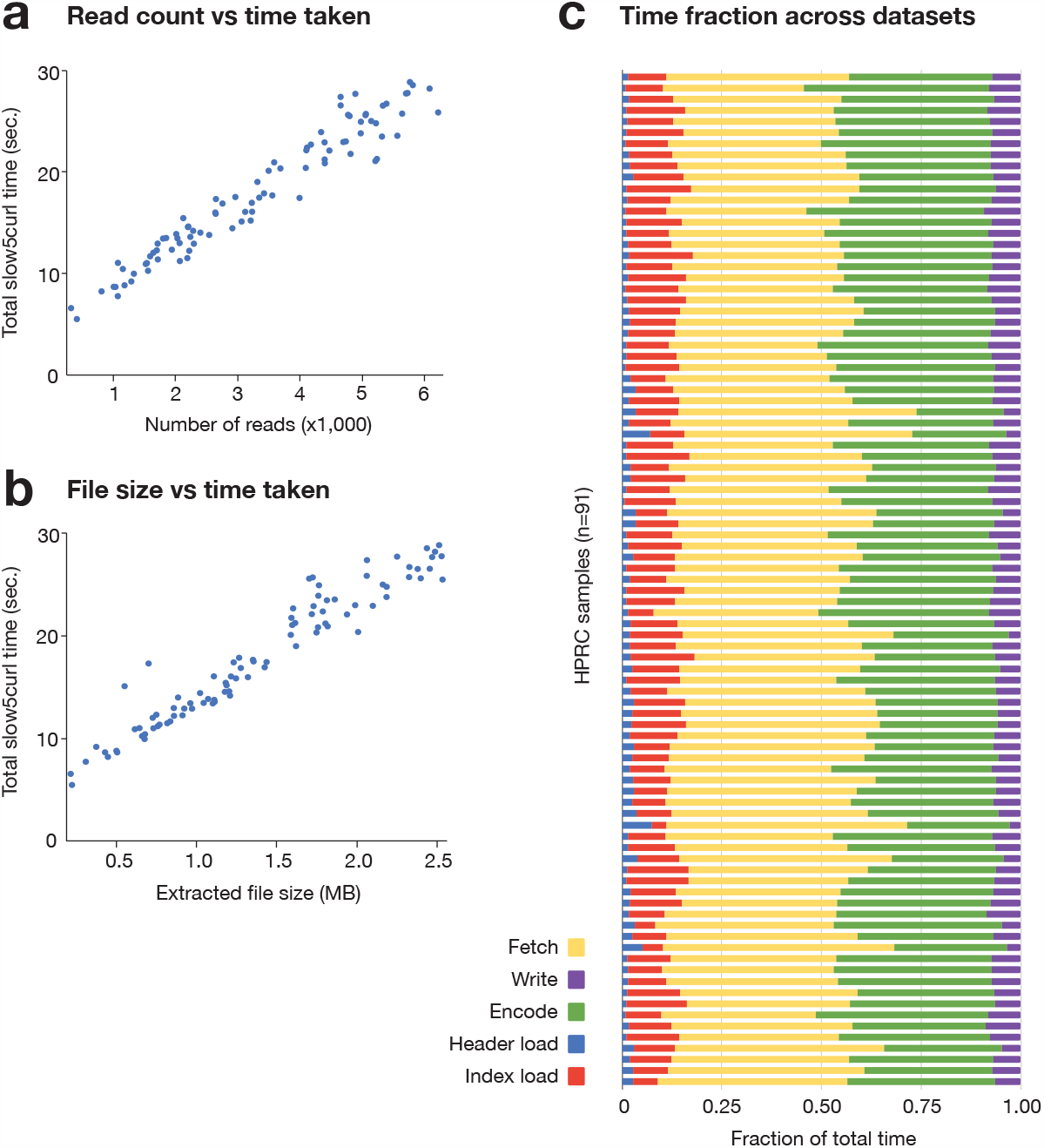
Evaluating slow5curl performance on large cohort datasets. (**a**) Time taken to fetch all signal reads corresponding to a hypothetical gene panel (n = 100 genes) from each of n = 91 whole-genome ONT sequencing datasets currently available via the Human Pangenome Reference Consortium (HPRC), relative to the number of signal reads being extracted for each dataset, which varies depending on the sequencing depth for each HPRC sample. (**b**) Same as above but fetch times are shown relative to extracted file sizes (in MBytes). The linear correlation observed in these two plots indicates slow5curl maintained stable rate of data read fetching across the full HPRC cohort. (**c**) Stacked bar chart shows the fraction of total time taken to fetch reads from each HPRC sample allocated to each individual component of the process (‘fetch’, ‘write’, ‘encode’, ‘header load’, ‘index load’).

